# Reproductive biology of three *Ziziphus* species (Rhamnaceae)

**DOI:** 10.1101/077289

**Authors:** M. Kulkarni

## Abstract

The reproductive biology of three *Ziziphus* species: *Z. jujube*, *Z. mauritiana* and *Z. spina-christi*, with respect to seasonal growth stages, floral biology, pollen performance, fruit set and seed viability were compared. To elucidate the reasons for variations in reproductive success we also estimated the ploidy levels among the genotypes of these tree species. Four genotypes of *Z. mauritiana* and *Z. jujube* each, and two natural selections of *Z. spina-christi* were grown in the Negev highlands at the Sde Boqer experimental plot. Field observations were conducted to determine seasonal growth stages and floral biology; laboratory studies included florescent microscopy, differential interference contrast microscopy, *in vitro* pollen germination and histological techniques. Seasonal growth stages were similar in *Z. spina-christi* and *Z. mauritiana* but distinct in *Z. jujube*, which was the earliest species. Flower opening stages were similar in all species and genotypes. Anthesis timing was genotype specific and did not depend on species. Anthesis types were subdivided into early and late morning as well as afternoon. We found a wide range of values for the number of flowers per inflorescence between species and genotypes, but all genotypes set one fruit per inflorescence with the exception of *Z. jujube* genotype Tamar, which set two fruit per inflorescence. Significant variation in fruit weight was observed between species as well as genotypes. Fruit of *Z. mauritiana* and *Z jujube* weighed between 9 to 37 g while fruit weight of *Z. spina-christi* was between 0.5 to 0.7 g. Flowers of all species and genotypes had two ovules but viable seed set per fruit differed dramatically between species under open pollination. Viable seed set was about 18% for *Z. mauritiana* and *Z jujube*, while *Z. spina-christi*, had the highest viable seed set at about 84%. Histological analysis revealed post fertilization embryo abortion which may be responsible for the relatively low reproductive success in *Z. mauritiana* and *Z. jujube*. The results obtained in this study provide an important basis for selecting elite complementary genotypes toward the optimization of *Ziziphus* fruit production in semi-arid regions.

## Introduction

*Ziziphus* (Rhamnaceae) comprises about 170 species of spiny shrubs and small trees distributed in the warm-temperate and subtropical regions throughout the world (Islam, M.B. 2006). Three species were chosen for this study, two fruit crops: *Z. jujube* native to China (Saran, P.L. 2006)and *Z. mauritiana* native to India (Kumar Shukla, A. 2004) and *Z. spina-christi*, a non-commercial species endemic to the Middle East whose fruit, leaves and wood are exploited by native populations (Saied, A.S. 2008). *Z. mauritiana* is adapted to hot arid regions (Mizrahi, Y. 1996) while *Z. jujube* is adapted to both hot and cold climates, capable of surviving periods well below freezing (Azam-Ali, S.N. 2006). *Z. spina-christi* also has a wide temperature tolerance range growing in Mediterranean drylands with temperatures as low as −2° C (Azam-Ali, S.N. 2006).

*Z. jujube* is deciduous tree, flowering from April to June and fruit matures August to October in China. *Z. mauritiana* is an evergreen tree, and flowers from August to October, its fruit mature from January to March in India (Kumar Shukla, A. 2004). *Z. spina-christi* is evergreen but may lose its leaves during cold winters. It flowers from May to December and fruit matures in spring (Galil, J. 1967). Phenological response of these two newly introduced species under Israeli condition needs to be checked for studying fruit set and fruit growth in comparison to their native grown countries.

Flowers of *Ziziphus* are hypanthium type and inflorescences are cyme or small panicle (Vashishtha, B.B. 1979). The flowers are small and inconspicuous; their most noticeable part is a pale yellow calyx which is 6-8 mm in diameter. Prominent pollinating insects are honey bees, yellow wasps and other hymenopterous species (Kumar Shukla, A. 2004). The life of an individual flower is very short (2-3 days) and many flowers remain unpollinated in an inflorescence, eventually dropping down. The number of flowers per inflorescence range from 12-16 in *Z. mauritiana* (Vashishtha, B.B. 1979;Teaotia, S.S.) with a similar range in *Z. jujube* (Liu, Ping 2004; Liu et al.; 2008a) reported abundant flower production, high flower drop (87.9 to 99.9 %) with very low fruit set (1.1%) in *Z. jujube*.

*Ziziphus* flowers are synchronous protandrous dichogamous, i.e., pollen availability precedes stigma receptivity with little overlap between the sexual stages. Morning and afternoon type was studied in *Z. mauritiana* (Josan, J.S. 1980; Desai, U.T. 1986; Tel-Zur, N. 2008) and *Z. spina-christi* (Galil, J. 1967). Using suitable complementary morning and afternoon type cultivars is vital for optimum fruit production as was reported in other dichogamous species as avocado (*Persea americana* Mill.) (Gazit, S. 2002).

The existence of a range of self and cross incompatibilities in *Ziziphus* species is known (Pareek, O.P. 1996). Self incompatibility as well as very low level of cross-compatibility was reported in *Ziziphus celata* (Weekley, C.W. 2002). Self incompatibility was reported by Galil and Zeroni (Galil, J. 1967) in *Z. spina-christi*. Self as well as cross incompatibility is also reported in *Z. mauritiana* cultivars (Teaotia, S.S.; Godara, N.R. 1980; Faroda, A.S. 1996). Interestingly, Liu et al. (2008) reported 87.8% out of 180 *Z. jujube* cultivars set fruits under self pollination.

The objective of this study was to compare reproductive biology of three *Ziziphus* species: *Z. jujube*, *Z. mauritiana* and *Z. spina-christi*, with respect to seasonal growth stages, floral biology, pollen performance, and fruit set and seed viability. To elucidate the reasons for variations in reproductive success we also estimated the ploidy levels among the genotypes of these tree species. This information will be further used to enhance fruit production, and to improve breeding programs.

## Materials and methods

### Study site and plant material

This study was conducted in an experimental plot at the Sede-Boqer Campus of Ben-Gurion University of the Negev, located in the Negev Highlands, Israel (30°52” N, 34°46” E, 430 m above sea level) during April 2007 to December 2008. The Negev highlands are characterized by cold and mostly sunny winters, with mean daily maximum/minimum temperatures of 14.9/3.8°C, and by hot, dry summers, with mean daily maximum/minimum temperatures of 32/17°C (Fig.1). Average annual rainfall is 80 mm, with considerable deviation from year to year. Data for mean temperature and relative humidity was obtained from meteorological station at the Dept. of Solar Energy and Environmental Physics, BIDR, BGU, Sede Boker Campus.

**Fig. 1.**
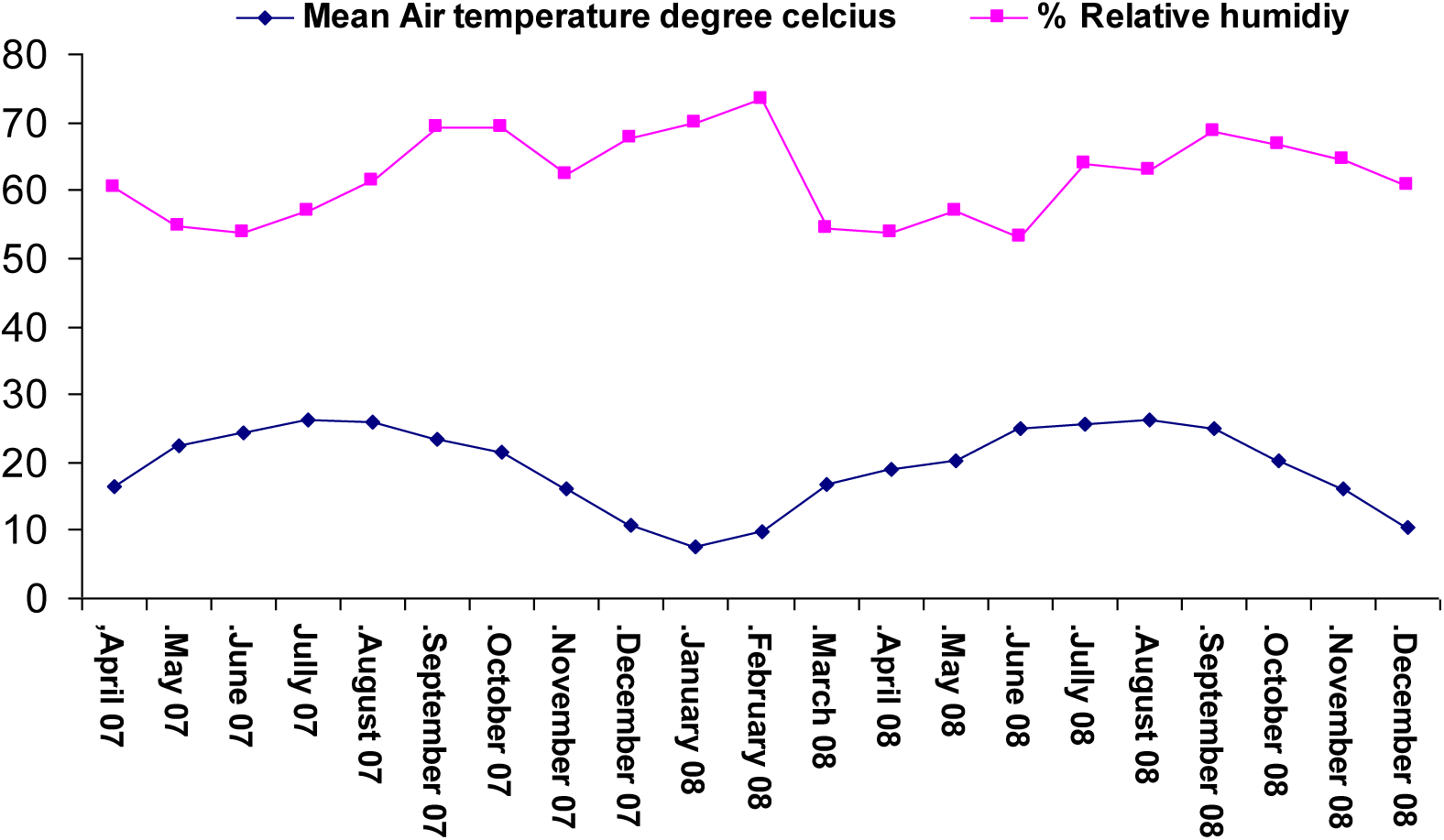
Mean air temperature and relative humidity during experimental period

Four *Z. jujube* genotypes (Lang, Tamar, Ben Li and Li); four *Z. mauritiana* genotypes (Seb, Q-29, Umran and B5/4) and two *Z. spina-christi* genotypes (Bachara and Or HaNer), were used in the present study. *Z. jujube* and *Z. mauritiana* genotypes were grafted on suitable rootstock whereas *Z. spina-christi* genotypes were seedlings. The trees were 3 years old at the time of the study and were grown in a field with sandy loam soil. Chemical fertigation (Poly-Feed DRIP 23:7:23+2MgO with micronutrients, Haifa Chemicals Ltd.) was supplied through a drip system. The plants were irrigated with 56 L/week during the hot season and 16 L/week during the cold wet season (November-March).

### Seasonal growth stages

Five phenological stages were monitored every two weeks during April 2007 to December 2008: 1) dormancy break, i.e., beginning of vegetative growth, 2) initiation of flowering, 3) peak flowering, 4) fruit growth and maturation, 5) leaf shed, i.e. beginning of winter dormancy.

### Floral biology, fruit set and size

Flower opening phases was studied in 50 flower buds per tree. The flowers were randomly selected and tagged one day before anthesis and flower opening phase was monitored at 2-hour intervals over a one-month period. Phase categories were based on Galil and Zeroni (1967). Number of flowers per inflorescence was studied in 40 inflorescences. The inflorescences were randomly selected and tagged for each tree; the number of flowers per inflorescence was recorded. Onset of anthesis was observed in 50 randomly selected inflorescences. The inflorescences were tagged; time of anthesis was monitored at various times of the day over a four-week period. Stigma receptivity was checked by fixing at least 30 stigmas at each flower opening stage as described by Barcelo et al. (2002). Images were viewed with an Axioimagera1 LED (Zeiss) microscope and captured with digital camera [Zeiss Axiocam HRC].

Mature fruit was collected, weighed and seed set, under open pollination conditions, was determined in all genotypes as a measure of their comparative reproductive success.

### Pollen performance

To determine pollen viability and diameter ten flowers per genotype were collected at anthesis. Pollen grains from undehisced anthers (two anthers per flower) were placed in 0.5 M sucrose solution in a 1.5 ml eppendorf tube, vortexed and dispersed on a microscope slide in a drop of FDA stain (2 μg/mL), incubated for 4-5 min at room temperature in the dark, and examined with an Axioimagera1 LED (Zeiss) fluorescent microscope. Brightly fluorescing grains were scored as viable, while weakly fluorescing grains or those not exhibiting fluorescence were scored as non-viable. At least 500 grains were examined per flower. In addition, the equatorial diameter of 100 viable pollen grains was measured using AxioVision AC Rel. 4.5 program (Zeiss).

*In vitro* pollen germination tests were carried out as described by Tel-Zur and Schneider (Tel-Zur, N. 2009). Twenty flowers (five from each genotype) were sampled and anthers were removed, placed in 0.5 M sucrose solution, vortexed and dispersed on germination medium consisting of 1% (w/v) agarose, 10% (w/v) sucrose, 300 mg/L of Ca(NO_3_)_2_ and 100 mg/L of H_3_BO_3_ on microscope slides. Slides were incubated at 25°C under dark condition for 24 h and pollen germination was observed using an Axioimagera1 LED (Zeiss) microscope. Pollen grains were classified as germinating when the pollen tube length exceeded the pollen diameter. Percent pollen germination was calculated from the germination rates of three replications of 300 grains.

*In vivo* pollen tube growth was determined in at least 20 flowers in each of the four *Z. jujube* genotypes. Approximately 12 h prior to anthesis flowers were selected, tagged and emasculated. Self pollination was performed approximately 4 hrs after anthesis when the stigma became fully receptive. Flowers were collected at anthesis and stored for four hours at 4°C until used for pollination. The styles were collected at the following time intervals: 4, 8, 16, and 24 h after pollination, and *in vivo* pollen tube growth was observed using the aniline blue protocol according to Mori et al. (2006). Pollen tubes were observed with an Axioimagera1 LED (Zeiss) fluorescent microscope and pictures were taken with Zeiss Axiocam HRC. AxioVision AC Rel. 4.5 program (Zeiss) was used to measure pollen tube growth from the tip of the stigma. Pollen tube growth rate was estimated by measuring pollen length at defined time intervals following pollination. The initial number of pollen grains germinating on the stigma and the number of pollen tubes continuing growth were also determined.

### Seed viability

Seeds were removed from mature fruit. The number of viable (fully developed) and aborted (collapsed, shrunken) embryos, was recorded. Viable as well as aborted seed weight was determined. Germination tests for seed viability testing was carried out by soaking seeds for 48 h in water and checking germination in Vermiculite 2 (Agrekal, Habonim Industries, Moshav Habonim, D.N. Hof HaCarmel 30845, Israel). Twenty seeds per genotype were tested for germination in two replications.

Anatomy of viable and unviable seed was studied using histological analysis, as described by (Tel-Zur, N. 2009). UV fluorescence was used to study autofluorescence of nuclei and protein bodies using an Axioimagera1 LED (Zeiss) fluorescent microscope. Photographs were taken with a Zeiss Axiocam HRC and captured using AxioVision AC Rel. 4.5 software program (Zeiss).

### Female gametophyte studies

Twenty flowers were collected from all genotypes of *Z. jujube* to identify abnormalities which may explain embryo abortion. Flowers were collected 4 hrs after anthesis, their ovules were removed, treated with cell wall degrading enzyme mixture and cleared as described by Wei and Sun (2002). Cleared ovules were examined using Nikon Eclipse E 600 microscope equipped with differential interference contrast microscopy (DIC) optics.

### Flow cytometry

Flow cytometry analysis was performed as described by Garcia et al. (2009). *Pisum sativum* cv. Citrad was used as standard with known genome size of 9.39 pg (Johnston, J.S. 1999). Genome size of *Ziziphus* genotypes was calculated according to Dolezel and Bartos (Dolezel, J. 2005). *Z. mauritiana* cv. Umran was reported to be tetraploid based on chromosome count (2n=4x=48) by Gupta et al. (2003) thus was used to estimate ploidy level of the rest of the genotypes.

### Statistical analysis

Differences between genotypes for measured parameters were tested for significance using one way ANOVA and mean separation with the Tukey HSD-test, with confidence limits set at p<0.05 using JMP 5.0.1 statistical analysis software. Arcsine transformation to normalize data was done for percentages. Seed viability percentage was analyzed at species level by pooling data from all genotypes from each species.

## Results

### Seasonal growth stages

Five seasonal growth stages were identified in all the three species: 1) winter dormancy, 2) vegetative growth, 3) early flowering, 4) full bloom, 5) fruiting. Phenology varies slightly between *Z. mauritiana* and *Z. spina-christi* but it was distinct in *Z. jujube* (Fig. 2).

**Fig. 2.**
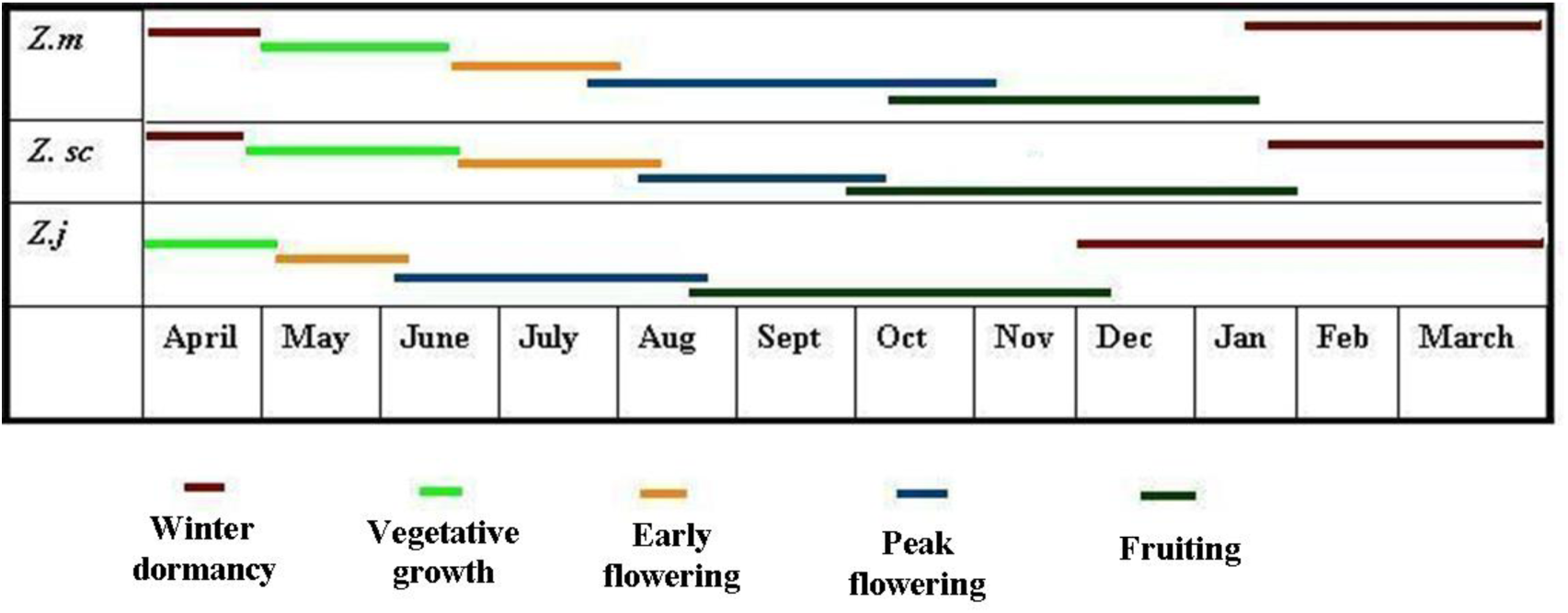
Comparative phenophases of *Ziziphus* species. *Z. jujube* (*Z. j*), *Z. mauritiana* (*Z. m*) and *Z. spina-christi* (*Z. sc*) under prevailing growing conditions in the Negev desert of Israel.

Fruiting period for *Z. jujube* is from Mid August to December depending on genotype. Tamar and Ben Li had earlier flowering (first week of April) and fruiting than Li and Lang (third week of April), with the longest flowering and fruiting periods in Tamar (April to end of July and Mid June to September). In general, *Z. jujube* had earlier flowering and fruiting stages as compared to the *Z. mauritiana* and *Z. spina-christi* genotypes. In the latter two species, peak flowering period was between Jully to November, fruiting period between October to January, and winter dormancy between February to May (Fig.2).

### Floral biology, fruit set and size

Flower opening was divided into six different stages in all genotypes, similar to that described by Galil and Zeroni (1967) (Fig. 3 A to F). Stigmata are not receptive at the time of anthesis (Fig. 3 G, H) and they become receptive 4 hrs after anthesis in all genotypes (Fig. 3 I, J). Stigmata begin losing receptivity 16 hrs after anthesis and become non-receptive 24 hrs after anthesis (Fig. 3 K, L). Anther dehiscence in all genotypes starts at anthesis and continues for 2 hrs (Fig. 3 N, O). Anthers are devoid of pollen grains at 4-5 hrs after anthesis (Fig. 3 P, Q and R). Flower opening stages were similar in all species and genotypes.

**Fig. 3.**
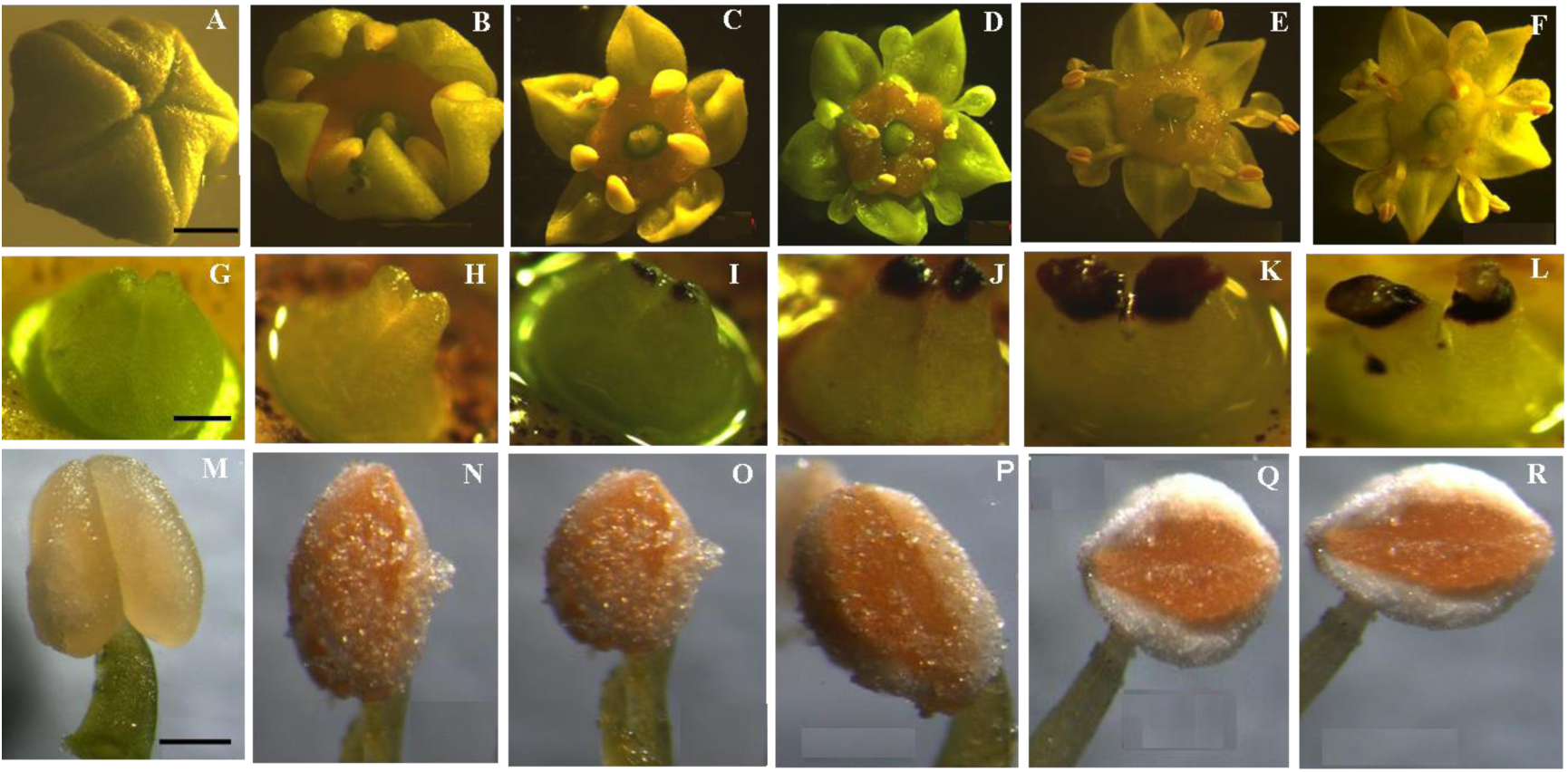
**a-f** Flower opening phases in *Z. jujube* genotype Li. **a** Before anthesis. **b** 2 hrs after anthesis, slits appear between petals and flower opens gradually. **c** 4 hrs after anthesis, flower is wide open, stamens are erect, enveloped by petals. Style is short and stigmata undeveloped. **d** 8 hrs after anthesis, petals gradually diverge, stamens devoid of pollen. **e** 16 hrs after anthesis, petals and stamens recurve between sepals. **f** 24 hrs after anthesis, floral structure same as **e** but anthers are dry. **g-l** Stages of stigma receptivity. **g** 2 hrs before anthesis, stigmata non-receptive and undeveloped. **h** 2 hrs after anthesis, stigmata non-receptive and undeveloped. **i** 4 hrs after anthesis, stigmata start developing. **j** 8 hrs after anthesis, stigmata fully receptive. **k** 16 hrs after anthesis, stigmata start loosing receptivity and are fully developed. **l** 24 hrs after anthesis, stigmata non-receptive and start withering. **m-r** Stages of pollen dehiscence. **m** At time of anthesis. **n** Peak dehiscence at 2 hrs after anthesis. **o** Continued dehiscence at 4 hrs after anthesis. **p** Dehiscence almost complete 6 hrs after anthesis and anthers devoid of pollen grains. **q-r** 8 to 10 hrs after anthesis, anthers empty. *Scale bar* 200 μm in **g-r**; and 1 mm in **a-f**.

Genotypes could be divided into three groups according to timing of anthesis onset, i.e., early morning [06:00-08:00] type, late morning [08:00-10:00] type and afternoon [12:00-14:00] type. Morning types are in the “male phase” in morning and in “female phase” in afternoon, whereas afternoon types are in the “male phase” in the afternoon and female phase in the “evening” (Table 1).

The number of flowers per inflorescence ranged from 7.5 to 23.7 in the three species studied and differed significantly between genotypes. *Z. jujube* genotype Li had the fewest whereas Tamar had the greatest number of flowers per inflorescence (Table 1; Fig. 4a,b). *Z. spina-christi* genotypes Bachara and Or HaNer had 15.6 to 16.4 flowers per inflorescence (Table 1; Fig 4c). The number of flowers per inflorescence in *Z. mauritiana* was dependent on genotype, Umran having and average of 9.4 and Seb with 17.7 (Table 1; Fig. 4d).

**Fig. 4.**
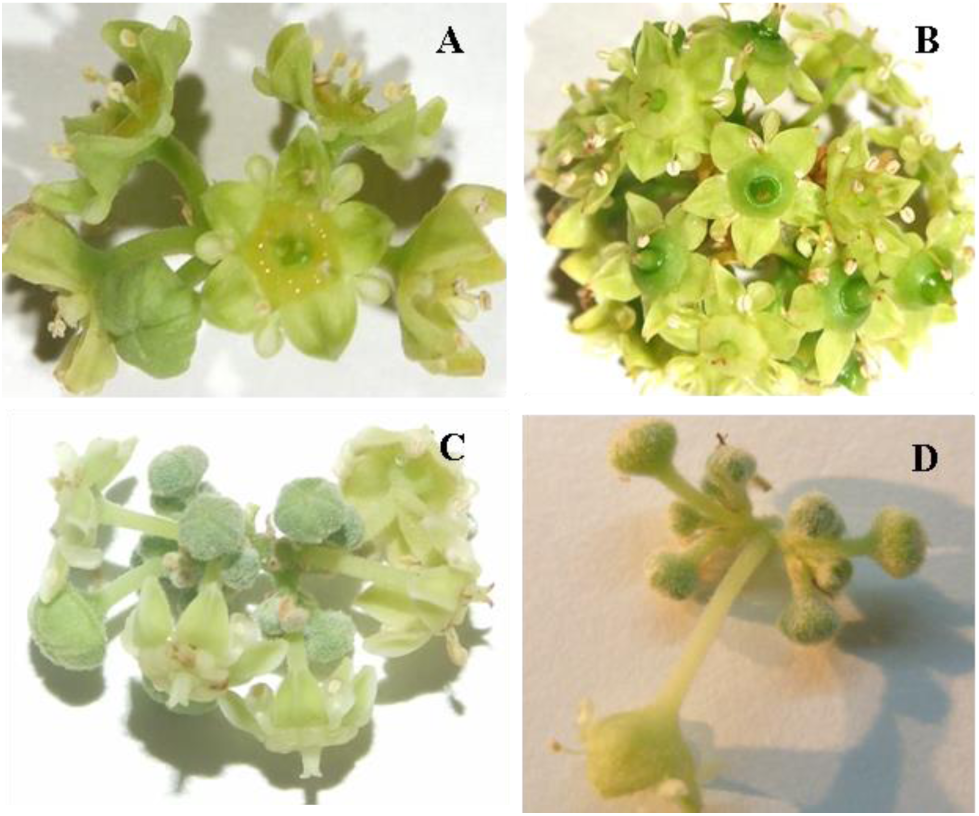
Variation in number of flowers per inflorescence in *Ziziphus* genotypes. **a** *Z. jujube* genotype Li. **b** *Z. jujube* genotype Tamar. **c** *Z. spina-christi* genotype Or HaNer. **d** *Z. mauritiana* genotype Umran.

The number of fruit set per inflorescence was determined ca. 4 weeks after open pollination. Although *Ziziphus* spp. produced numerous flowers on each inflorescence, fruit set was very low irrespective of species or genotypes (Table 1). One fruit per inflorescence was recorded on all genotypes with the exception of *Z. jujube* genotype Tamar, which set two fruits per inflorescence.

Significant genetic variation in fruit weight was observed between species as well as genotypes (Table 1). *Z. jujube* genotype Li had the largest fruit among all genotypes studied (37.3 g). Lang and Ben Li had medium size fruits ranging from 17.9-24.2 g. Tamar had the smallest fruit among *Z. jujube* genotypes with an average weight of 9.2 g. *Z. mauritiana*, genotypes Umran and B 5/4 had significantly bigger fruits with average weights of 33.6 and 29.5 g, respectively. Seb and Q-29 had medium fruits with average weights of 21.6 and 19.6 g, respectively. *Z. spina-christi* genotypes had very small fruits as compared to the two cultivated fruit crop species. Average fruit weights of Bachara and Or HaNer were only 0.51 and 0.74 g, respectively.

### Pollen performance

The mean pollen grain diameter for each genotype is presented in Table 2. *Z. mauritiana* genotype B5/4 had smallest pollen grain diameter (23.7 μm) whereas *Z. spina-christi* genotype Bachara had largest pollen grain diameter (28.2 μm). Mean pollen diameter for *Z. mauritiana* and *Z. jujube* genotypes was 26.1 μm and 25.3 μm respectively while it was significantly (p<0.05) higher (28.1 μm) in *Z. spina-christi*.

The percentage of viable pollen as determined with FDA was between 60.6 to 80.4% and differed significantly between species (Table 2; Fig. 5a). Mean pollen viability for all genotypes of *Z. mauritiana* and *Z. jujube* was 66.4% and 68.0 % respectively, and significantly higher (79.2%) for *Z. spina-christi*. The lowest pollen viability was observed in *Z. mauritiana* genotype Umran (60.6%) whereas the highest pollen viability was observed in *Z. spina-christi* genotype Or HaNer (80.4%).

**Fig. 5.**
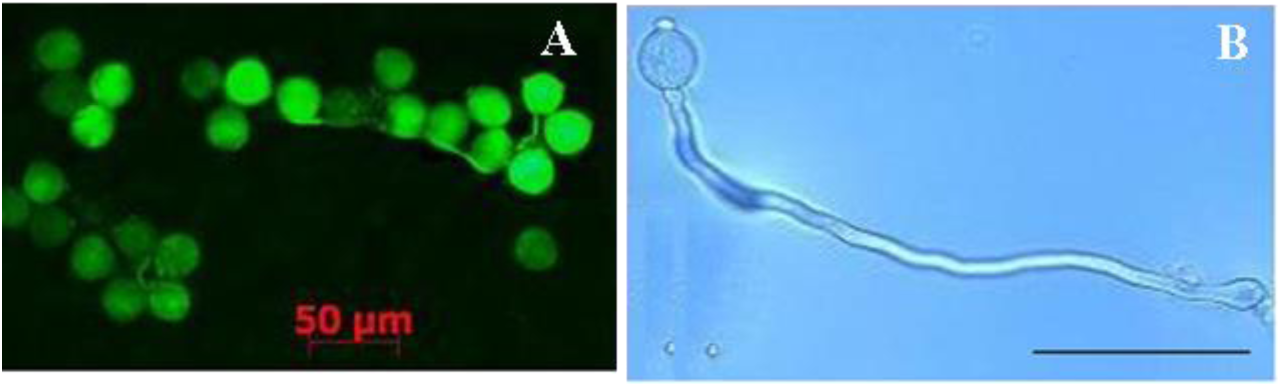
Pollen performance in *Z. jujube* genotype Li. **a** Pollen viability test using FDA stain, fluorescing grains are viable. **b** *In vitro* pollen germination, 12 hrs after plating on germination medium. *Scale bar* 50 μm in **a**; and 100 μm in **b**.

Significant differences for *in vitro* pollen germination rate were observed between the genotypes of the three species under study (Table 2; Fig. 5b). Mean *in vitro* pollen germination for *Z. jujube* and *Z. mauritiana* genotypes, was 54.5% and 57.2% respectively and it was significantly higher (71.5%) in *Z. spina-christi*. Lowest pollen germination rate was observed in *Z. jujube* genotype Tamar (49.6%) whereas highest *in vitro* pollen germination was observed in *Z. spina-christi* genotype Or HaNer (72.8%).

*In vivo* pollen tube growth studies following hand self-pollination were carried out for *Z. jujube*. Pollen tubes of all *Z. jujube* genotypes germinated normally and grew more than 500 μm in 24 hrs after pollination, reaching the base of the ovary (Fig. 6). Pollen tube growth kinetics was also studied on *Z. jujube* genotype Lang. Numerous pollen tubes initially grow down the style in bundles and only 1.2-1.5% of the pollen tubes reach the base of the ovary (Fig. 7).

**Fig. 6.**
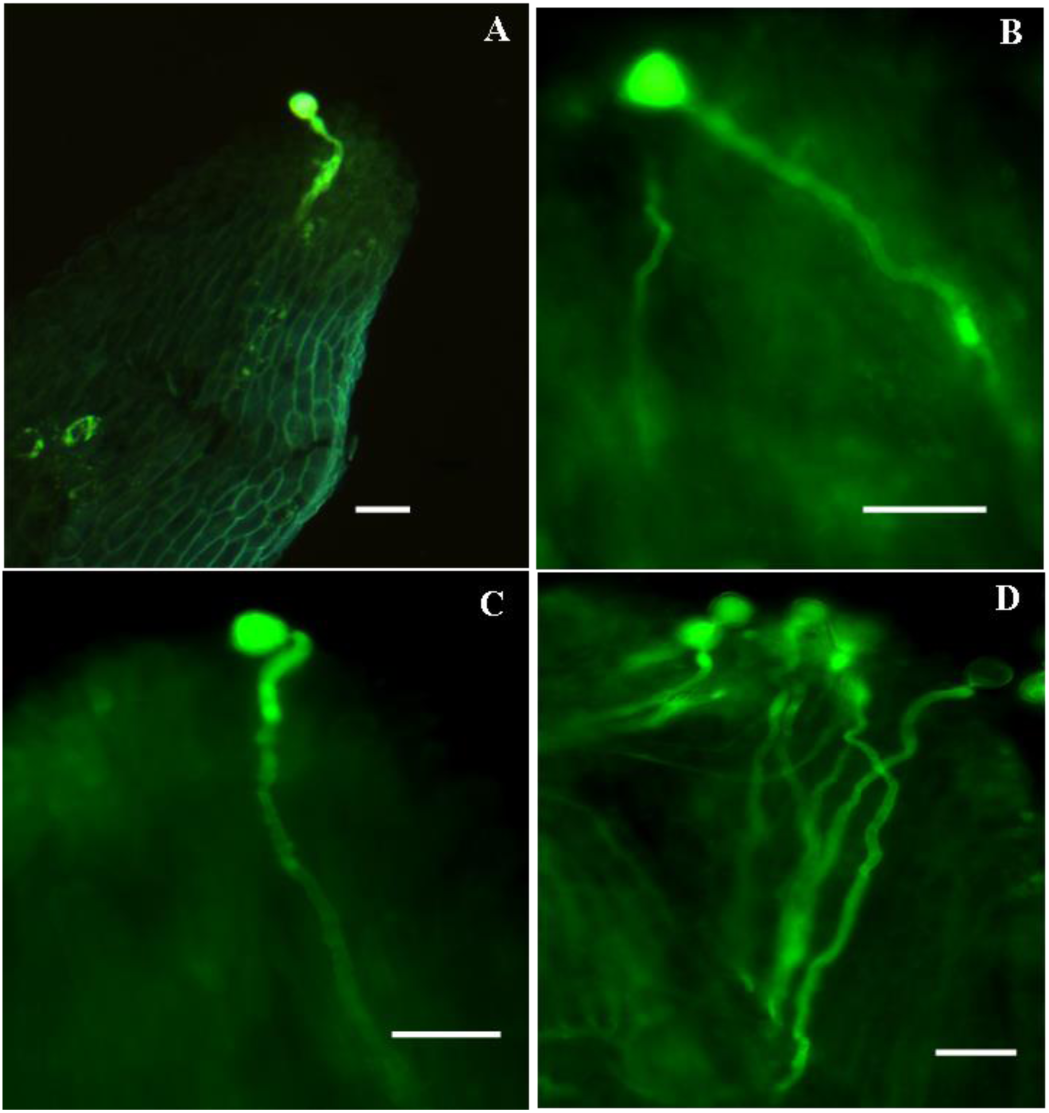
*In vivo* pollen tube growth in *Z. jujube* genotype Li after hand self-pollination. **a** Pollen tube growth after 4 hrs. **b** After 12 hrs. **c** After 16 hrs. **d** After 24 hrs. *Scale bar* 50 μm.

**Fig. 7.**
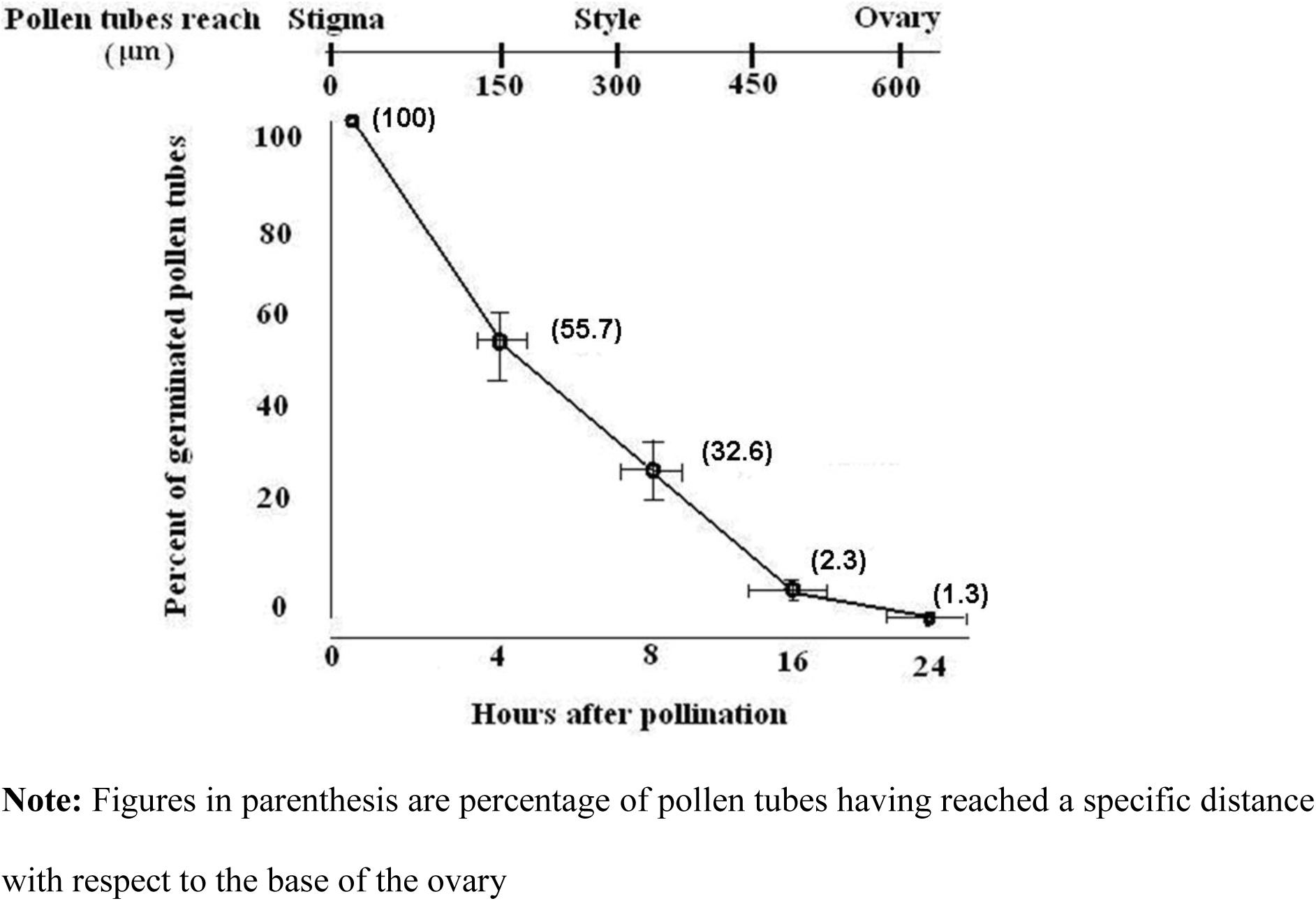
Kinetics of *in vivo* pollen tube growth in *Z. jujube* genotype Lang after hand selfpollination

The flowers were emasculated four hours after anthesis (peak of stigma receptivity)

**Note:** Figures in parenthesis are percentage of pollen tubes having reached a specific distance with respect to the base of the ovary

### Seed viability

All *Ziziphus* species and genotypes had two ovules in the ovary. Only one ovule set viable seed in *Z. mauritiana* and *Z. jujube*, the second embryo apparently aborted after fertilization (Fig. 8a). Among the 830 fruits opened in these two species, none of the fruits was with two viable seeds. *Z. mauritiana* and *Z. jujube* genotype fruits were either single seeded or without any viable seeds. A single viable seed was observed in 19.7% of fruits in *Z. jujube* and 17.5% of fruits in *Z. mauritiana* (Table 3). The remaining fruits did not have any viable seeds and an aborted embryo at different stages was observed. In *Z. spina-christi*, both ovules were fertilized maturing into viable seeds in more than 84.2% of stones (Table 3; Fig. 8b).

**Fig. 8.**
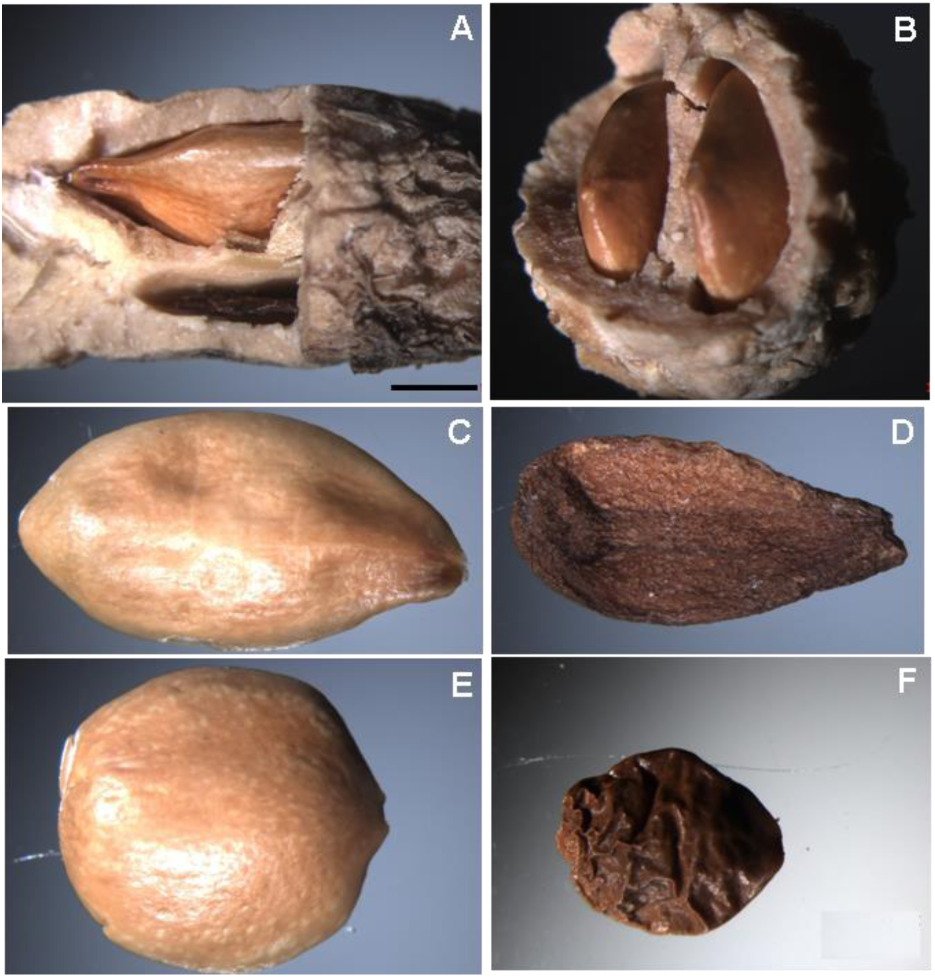
Reproductive success in *Z. mauritiana* genotype Seb as compared with *Z. spina-christi* genotype Or HaNer. **a** Only one seed is fully developed and viable in stones of *Z. mauritiana* whereas the second seed is aborted. **b** Two fully developed, viable seeds in stone of *Z. spina-christi*. **c** Fully developed viable seed of *Z. mauritiana*. **d** Aborted, unviable seed of *Z. mauritiana.* **e** Fully developed viable seed of *Z. spina-christi*. **f** Aborted, unviable seed of *Z. spina-christi. Scale bar* 1 mm.

Viable seeds (Fig. 8c, e) of all species had significantly higher average seed weight as compared to aborted seeds (Fig. 8d,f). *Z. spina-christi* fruit size is very small but seed size was similar to the other two species (Table 3). Viable seeds of all species had a high germination rate ranging between 61.3 to 83.7 (Table 3). Seeds germinated within one week and developed into normal healthy seedlings after one month. None of aborted seeds imbibed water and germinated.

Histological comparison of viable and aborted seeds revealed remarkable differences at the anatomical level. Average seed thickness of normal seed (Fig.7) was about twice that of aborted seeds (Fig. 8). Cotyledons of normally developing seeds had normal xylem vessel development (Fig 9a) whereas these structures are totally absent in aborted seeds (Fig. 9b). Exotesta and hourglass cell layers were thicker in viable seeds, with an average thickness 42.5 and 40.3 μm, respectively; whereas these layers were collapsed and underdeveloped in aborted seeds with an average thickness 26.3 and 11.5 μm, respectively (Fig.9c-f). Cotyledon cells had normal cytoplasm, protein bodies and nucleus in viable seeds (Fig. 10a, c, e) where as cotyledon cells in aborted seeds were extremely irregular in size and shape with virtually no nuclei and cell content are visible indicating dead cells (Fig.10 b,d,f).

**Fig. 9.**
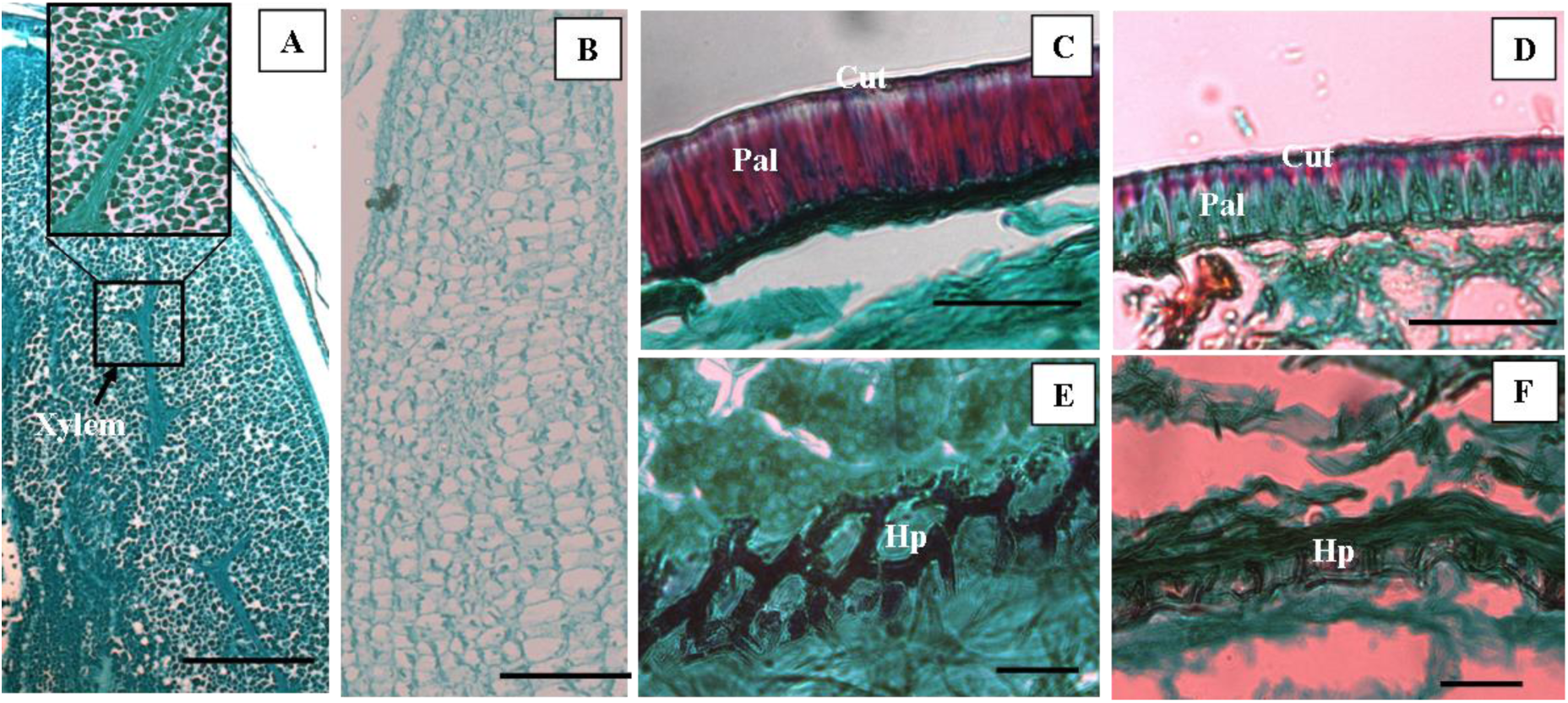
Histological studies of viable and aborted seeds in *Z. jujube* genotype Li. **a** Cotyledon of viable seeds. Enlarged square showing normal xylem vessels. **b** Shrunken cotyledon in aborted seeds without xylem vessels. **c** Thicker exotesta in normal seed. **d** Shrunken exotesta in aborted seeds. **e** Thicker hourglass-cell layer in normal seed. **f** Thinner and collapsed hourglass-cell layer in aborted seed. *Scale bar* in **e, f** - 20 μm; in **c, d** - 50 μm; and in **a, b** - 100 μm. *Cut* cuticle, *Hp* hourglass cells of hypodermis, *Pal* palisade cells.

**Fig. 10.**
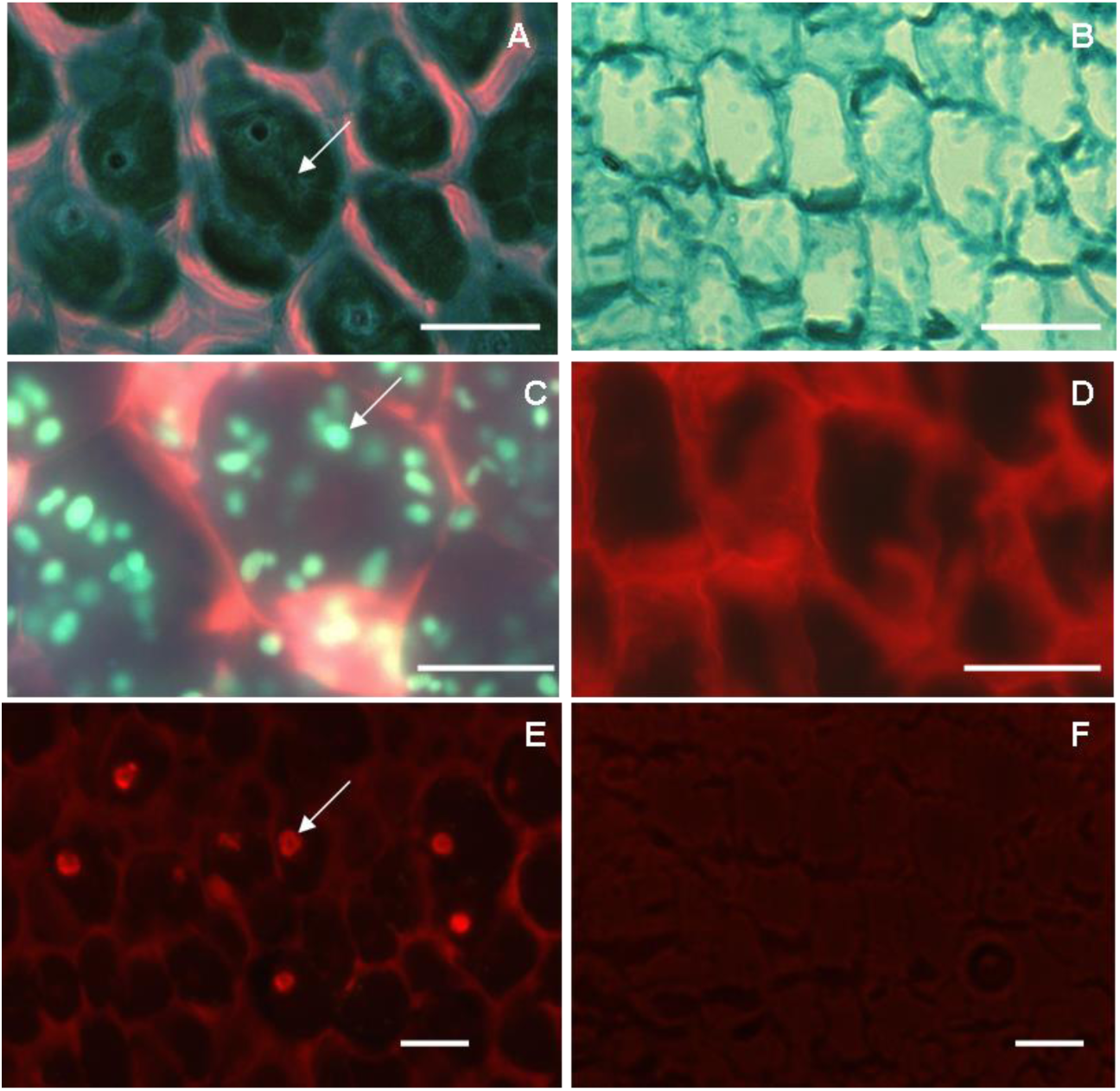
Histological observation of cotyledon cells in viable and aborted seeds in *Z. jujube* genotype Li. **a** Viable seed has fairly uniform cotyledon cells with well stained cytoplasm (arrow). **b** Aborted seed has irregular cotyledon cell size, devoid of cytoplasm. **c** Viable seed, showing a large number of protein bodies in cotyledon cells (arrow). **d** Aborted seed, lacking protein bodies in shrunken cotyledon cells. **e** Viable seed with well stained nuclei within cells of cotyledon (arrow). **f** Aborted seed cotyledon cells without stained nuclei. [**c-f** fluorescence excitation range 400-570 nm]. *Scale bar* 20 μm.

### Female gametophyte studies

An embryo sac was visible in each isolated ovule. The egg cell and central cells in the embryo sac become more visible after enzymatic treatment for 30 min at room temperature. DIC optics revealed a completely developed polygonum type embryo sac (Fig.11a, b).

**Fig. 11.**
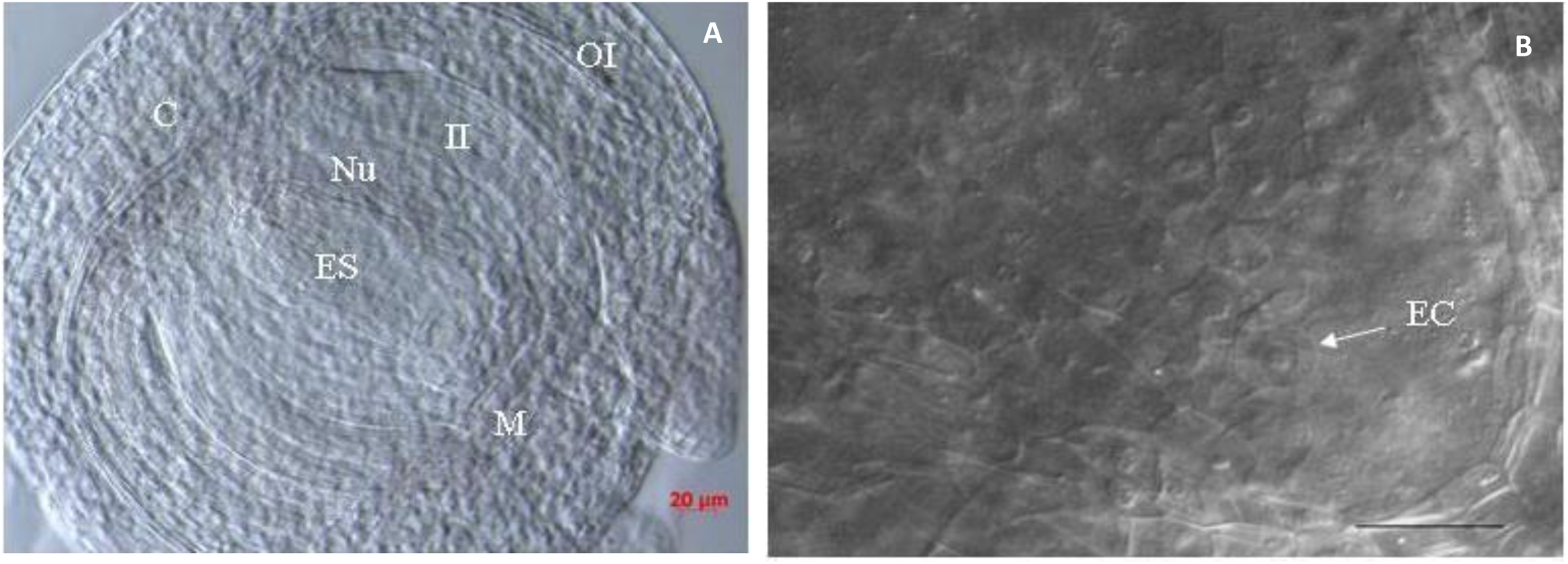
Ovule dissected from unpollinated ovary. (a) The dissected ovule was treated with enzyme solution, facilitating visualization of embryo sac. (b) Another dissected and enzymatically treated ovule. An egg cell is visible indicated by arrowhead. *Scale bar* 20 μm. *C* chalazal pole, *EC* egg cell, *ES* embryo sac, *II* inner integument, *M* micropylar pole, *Nu* nucellus, *OI* outer integument.

### Flow cytometry

Flow cytometry analysis indicated that all genotypes are diploid, with the exception of *Z. mauritiana* genotypes Seb and Umran which were tetraploid (Table 4; Fig. 12). Genome size calculated on the basis of peak values were 2.35-2.44 pg for *Z. mauritiana* genotypes Umran and Seb, as compared to 1.23 to 1.70 pg for the remaining genotypes (Table 4).

**Fig. 12.**
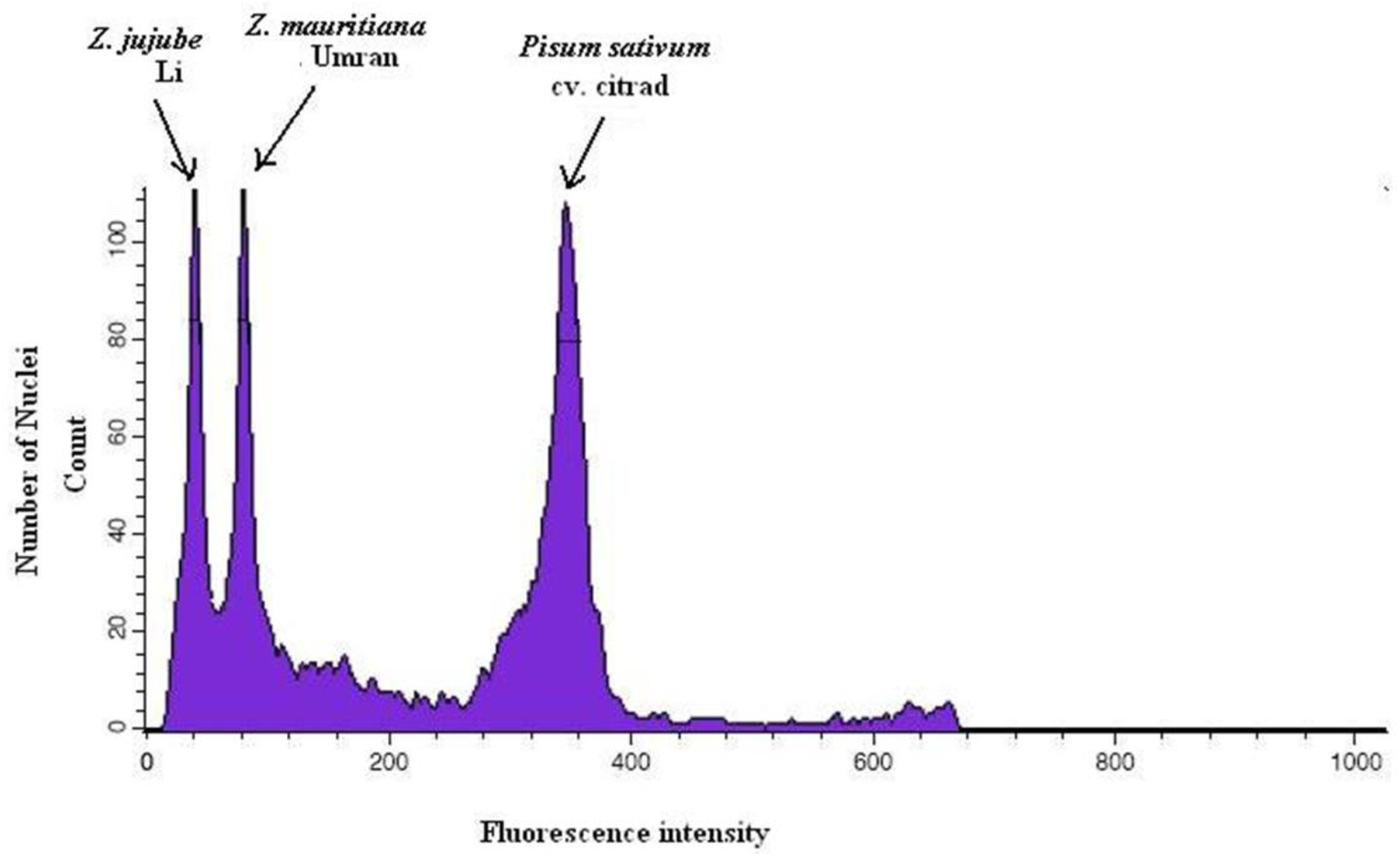
Representative nuclear DNA content histograms of *Z. jujube* genotype Li leaf nuclei. *Z. mauritiana* genotype Umran leaf nuclei sample was used as internal standard (tetraploid as reported by Gupta et al., 2003). *Pisum sativum* cv. Citrad was used for quantitative estimation of 2C nuclear DNA content.

## Discussion

### Seasonal growth stages

The three *Ziziphus* species in this study exhibited five distinct phenophases: 1) winter dormancy, 2) vegetative growth, 3) flowering, 4) full bloom, 5) fruiting. These stages are also described in their native countries (Kumar Shukla, A 2004). *Z. jujube* and *Z. mauritiana* genotypes exhibited distinct difference in their phenological phases studied in Negev desert conditions. These two species also behave distinctly for their phenophases as compared to their native countries. *Z. mauritiana* had flowering period between August to October and fruiting period between October to February under native conditions in India (Kumar Shukla, A 2004) but under Israeli conditions these cultivars had early flowering time (from Jully to October) and early fruiting (October to December). *Z. jujube* species had almost similar flowering and fruiting period as described in its native country i.e. China by (Kumar Shukla, A 2004). Dormancy period varied slightly, from December to April in Negev desert conditions of Israel whereas it is from November to March in China. Our studies indicated that *Z. mauritiana* as well as *Z. jujube* had at least one to one and half month earlier flowering as well as fruiting as compared to native countries which can be useful for commercial growers in Israel. It indicates possibility of early harvest and sell in to international markets as “out of the season” fruit crop.

*Z. jujube* being a frost tolerant species, sheds leaves early winter, and is the first species to break dormancy in spring; it also has the earliest flowering and fruit set. *Z. mauritiana* and *Z. spina-christi* had similar phenophase timing. Both are evergreen in temperate climates, however under cold desert conditions their leaves drop in winter. Phenological differences between *Z. jujube* and *Z. mauritiana* were also reported by (Desai, U.T. 1986; Babu, R.H. 1988; Sharma, V.P. 1990; Saran, P.L. 2005; Vishal, Nath 2002; Kim, W.S. 1982).

### Floral biology, fruit set and size

All three species and genotypes exhibited similar flower opening stages. The number of flowers per inflorescence in *Z. mauritiana* reported by (Vashishtha, B.B. 1979; Teaotia, S.S. 1964) 12-16 was similar to our findings (9.4 to17.7). Liu et al. (Liu, Ping 2004) reported a similar range of flowers per inflorescence in *Z. jujube* genotypes, however, we found greater variation among the various *Z. jujube* genotypes, ranging from 7.5 to 23.7.

Our studies indicate that, morning anthesis types under Negev desert conditions could be further sub-divided into early morning (6:00 to8:00 AM) and late morning (8:00 to 10:00 AM), see Table 1. Seb is early morning anthesis type genotype under Negev desert conditions, similar results anthesis time between 7:00 to 8:00 AM under Indian conditions is reported for Seb (Kumar Shukla, A. 2004). Umran is an afternoon anthesis type genotype under Indian conditions (anthesis 1:00 to 2:30 PM) as reported by (Kumar Shukla, A. 2004) but under Negev desert conditions it was categorized into late morning type (anthesis 8:00 to 10:00 AM) indicating changes in anthesis time under new growing conditions. Both *Z. spina-christi* genotypes in this study belonged to the afternoon anthesis type, although Galil and Zeroni (1967) reported morning as well as afternoon anthesis types. Little is known about *Z. jujube* genotype anthesis timings in published literature so this information from present investigation will be further helpful for optimizing fruit production of these genotypes under Negev desert conditions. The information gathered here will undoubtedly be helpful for further biological studies, utilizing complementary sexual morphs for optimum fruit production as reported by Gazit and Degani (1976) for avocado (*Persea americana* Mill.).

Although the number of flowers per inflorescence was variable between all genotypes, only one fruit set per inflorescence, with the exception of *Z. jujube* genotype Tamar, who set two fruits per inflorescence. This genotype also had the highest number of flowers per inflorescence (23.7). Liu et al. (2008) report that 91.5% of 180 genotypes in *Z. jujube,* bear only one fruit per inflorescence. This data indicates fruit set per inflorescence is not related to the number of flowers per inflorescence. Similar findings were reported by Reale et al. (2006) for various olive varieties, who further concluded that yield is related to the number of inflorescences per tree and not total flower production.

Considerable genetic variation in the genus *Ziziphus* exists for fruit weight was observed during present study. In this study fruit were round to ovate with fruit weight ranging from minimum 0.5 g for *Z. spina-christi* to maximum (37.3 g) for *Z. jujube* genotype Li. Fruit weight of the different genotypes and species grown at Sde Boqer is similar to that reported in previous work for *Z. mauritiana* (Saran, P.L. 2006; Ghosh, D.K. 2004) and *Z. jujube* (Mengjun, L. 2002) indicating normal fruit growth and development under presented environmental conditions. Wide variations of fruit shape and weight have been reported in the literature. Fruit weight ranged between 8 to 17g in *Z. mauritiana* (Saran, P.L. 2006; Ghosh, D.K. 2004) 2 to 46 g in *Z. jujube* (Mengjun, L. 2002) whereas *Z. spina-christi* fruits are comparatively very small weighing in range of 0.5 to 0.7g (unpublished data by authors). *Ziziphus* seeds are enclosed within a hard woody endocarp known as the stone. Stones vary in shape from round to subovate to ovate. Viable seeds per fruit ranged from one to two in *Z. mauritiana* under different agro-climatic conditions (Grice, A.C. 1996; Diallo, I. 2002; Danthu, P. 1992). Viable seed set was reported be very low in *Z. jujube* (Liu, P. 2008).

### Pollen performance

*Z. spina-christi* genotypes had the largest pollen diameter followed by *Z. mauritiana* and *Z. jujube* with the smallest. Desai et al. (1986) reported pollen diameter in the range of 26.9 to 30.3 μm for various *Z. mauritiana* genotypes and Mengjun (2002) reported 21.0 to 26.3 μm for several *Z. jujube* genotypes. Tel-Zur and Schneider, (2009) reported mean pollen diameter of 23.1 μm in B5/4 and 27.3 μm in Q-29 (both *Z. mauritiana*) which is consistent with the data presented here. The importance of pollen size lies in the fact that it can reflect the storage capacity for particular nutrients that may affect pollen tube growth (Roulston, T. 2000). *Z. spina-christi* being native species adapted to desert semiarid environment, and having significantly higher pollen diameter as compared to rest of to cultivated genotypes it could be assumed that large pollen grains have a competitive advantage over those with smaller grains under extreme environmental conditions. Should this hypothesis prove correct, then the pollen diameter trait may be used for selecting superior cultivars in breeding programs.

*Z. spina-christi* genotypes had the largest pollen diameter and the highest germination rates. Pollen size reflects the storage capacity for particular nutrients that affect pollen tube growth after germination (Roulston, T. 2000). Higher viability and pollen germination could be one of the contributing factors for higher reproductive success in *Z. spina-christi* as compared to *Z. mauritiana* and *Z. jujube*. Importance of greater pollen viability in donor parent to favor high fruit production is highlighted in other fruit crops. Orlandi et al. (Orlandi, F. 2005) report the importance of high pollen viability in olive (*Olea europaea* L.) production.

### Seed viability

*Ziziphus* seeds are enclosed within a hard woody endocarp known as the stone. Each fruit contains one stone embedded in the pulp at the centre of the fruit. Stones vary in shape and size from round to ovate and 1-2 cm. The number of seeds per fruit reported in previous studies ranged from one to two under different agroclimatic conditions in *Z. mauritiana* (Grice, A.C. 1996; Diallo, I. 2002; Danthu, P. 1992). Viable seed set is also reported be very low in *Z. jujube* (Liu, P. 2008), however, no published information exists about level of sexual reproductive success in *Z. spina-christi*.

In many species, low rates of seed reproduction are due to resource limitation (Stephenson, A.G. 1981), but resource limitation is an unlikely explanation for reproductive failure in *Z. mauritiana* and *Z. jujube* since it has been reported under a wide variation of environmental conditions (Liu, P. 2008; Grice, A.C. 1996). A wide range of reproductive success is represented in the three species studied here. At one end of this spectrum, *Z. spina-christi* is the most successful with a seed set of almost 6 times greater than the other two species *Z. mauritiana* and *Z. jujube*. Percentage viable seed set following open pollination varies significantly among the three species. It is highest in *Z. spina-christi* (84.2%) as compared to *Z. mauritiana* (19.7%) and *Z. jujube* (17.5%). Within species [between genotype] variation for seed set ranged between 11.0 to 22.1% in *Z. mauritiana* and 4 to 36% in *Z. jujube*. Post fertilization embryo abortion was observed in all the species studied here. In *Z. jujube* and *Z. mauritiana*, both ovules abort in more than 78% of fruits and only one viable seed is set in less than 18% of fruits (second locule with aborted seed). Grice (1996) also reported one viable seed per stone in *Z. mauritiana*. The reproductive success of *Z. spina-christi* is much higher; both ovules are fertilized and develop into normal viable seeds in more than 84% of fruits. The remaining 16% of the fruit are devoid of any viable seeds. A higher level of viable seed set in *Z. spina-christi* is reported by Saied et al. (2008). Liu et al (2008) reported viable seed set from 10 to 50% in 180 genotypes of *Z. jujube* under open pollination condition.

The process of post fertilization embryo abortion results in a low viable seed set in *Z. mauritiana* and *Z. jujube*. As pollen tube grows normally in all *Z. jujube* genotypes it could be hypothesized that *Z. jujube* genotypes are not self incompatible, at least at the sporophytic level. Although we did not follow process of fertilization and embryo development after selfing but histological analysis of viable as well as aborted seeds in all genotypes of three species indicate that under each case fertilization occurs and seed aborts only during development. Viable seed set under open pollination was studied since open pollination studies are a better indication of maximum reproductive potential, as a measure of reproductive success in comparison to controlled pollination conditions (Liu, P. 2008). This has been verified in other plant species (Medrano et al. 2000; Dhar et al. 2006). Self as well as cross incompatibility between various *Z. mauritiana* genotypes is well known (Pareek, O.P. 1996; Teaotia, S.S. 1964; Faroda, A.S. 1996) Self as well as very high level of cross incompatibility in *Ziziphus celata* i.e. Florida *Ziziphus* was also reported (Weekley, C.W. 2002).

Seed coat plays important role in control of water absorption, and hence on germination (Souza F.H.D.E. 2001). Wrinkled and collapsed seed coat in aborted seeds was impermeable as seeds do not imbibe and germinate at all. Thicker seed coat (almost double thickness) was observed in normally developed viable seed and was permeable as seeds imbibe and germinate. Similar to other flowering plants, in normally developed seeds storage products begin to accumulate after the cotyledon stage of embryo development. Significantly higher mature seed weight in normal seeds indicated accumulation of storage products in cotyledon cells whereas these storage products were absent in collapsed and dead cells of cotyledons in aborted seeds (as indicated by significantly lower mature seed weight). Similar results are reported by (Oliveira, D.M.T. 2005) while studying abortive seed development in *Ulmus minor*.

Another possible reason for abortive seed development could be abnormalities at female gametophyte level. Enzymatic digestion of unfertilized ovules with DIC optics revealed normal embryo sac development with with functional egg cell which excludes possibility of considering abnormality at female gametophyte level responsible for abortive seed development. As observed by histological analysis, in aborted seeds also embryo develops upto cotyledon stage indicating abortive seed development after fertilization only. In almost all abortive seeds, seed stops growing when embryo is at cotyledon stage, but at this stage all cotyledon cells are collapsed, dead without any storage products which are observed in normal seed cotyledon cells.

### Flow cytometry

Two ploidy levels were found in *Z. mauritiana* genotypes. Umran was reported as tetraploid (2n=4x=48) by Gupta et al. (2003) and we found similar genome size in Seb. The other two genotypes, Q-29 and B5/4 had genome size in range of 1.70 to 1.79 pg indicating their triploid genome size which could be responsible for lower reproductive success. It was interesting that *Z. jujube* genotypes also had lower reproductive success although all of them are diploid.

Reduced reproductive output of *Ziziphus* species may be due to incompatibility between maternal and paternal gametophytes. For example, two genotypes of *Z. mauritiana* were triploid and two genotypes were tetraploid. These ploidy level differences may lead to incompatibility leading to a very low level of reproductive success. Although, *Z. mauritiana* and *Z. spina-christi* flowered at similar times, presence of tetraploid *Z. mauritiana* genotypes (Seb and Umran) in the population did not affect the high levels of viable seed set in *Z. spina-christi*.

## Acknowledgement

We thank to J. Blaustein Center for Scientific Cooperation (BCSC), Ben-Gurion University of Negev for Postdoctoral fellowship granted to Dr. M. Kulkarni. The author thank Dr. Noemi Tel-Zur; R. Garcia, Dr. P. Zsolt and Mr. A. Cisneros for technical assistance. This work was partially supported by Jewish Fund for the Future-Goldinger foundation.

